# Composition and activity of the proteasome in human iPSC-derived neuronal model of early-stage sporadic Alzheimer’s disease

**DOI:** 10.64898/2026.06.23.734021

**Authors:** Aderemi Caleb Aladeokin, Michael Jeltsch, Hayk Davtyan, Mathew Blurton-Jones, Jari Koistinaho

## Abstract

**Introduction:** The proteasome is a critical cellular degradative machinery impaired in late-stage Alzheimer’s disease (AD). However, the status and activity of the proteasome in early-stage sporadic AD (sAD) is unknown.

**Methods:** A cellular model of human early-stage sAD was generated from sAD patient iPSC-derived cortical neurons by dual-SMAD inhibition. The iPSCs, neuroprogenitors, and cortical neurons were validated by the expressions of key markers. The level of total intraneuronal Aβ was measured by ELISA. Composition and native proteolytic activities of the proteasome in control and sAD cortical neurons were measured using complementary fluorogenic probes.

**Results:** Control and sAD patients iPSCs expressed pluripotent markers OCT4, NANOG, and SSEA4 which induced into neuroprogenitors expressing NESTIN and PAX6. The neuroprogenitors terminally differentiated into cortical neurons expressing neuronal markers MAP2 and TUJ1, and cortical layer marker TBR1. The level of intraneuronal Aβ in the sAD cortical neurons was significantly higher compared to control. Control and sAD cortical neurons expressed native 30S, 26S, and 20S proteasome assemblies with the sAD cortical neurons displaying higher 20S assemblies. Increased active 20S assemblies was associated with higher β1, β2, and β5 proteolytic sites activities.

**Discussion:** The significant elevation in the proteolytic activities of the β1, β2, and β5 subunits of 20S proteasome in sAD cortical neurons suggests that this may be a possible compensatory response to elevated intraneuronal Aβ.

**Graphical abstract:** 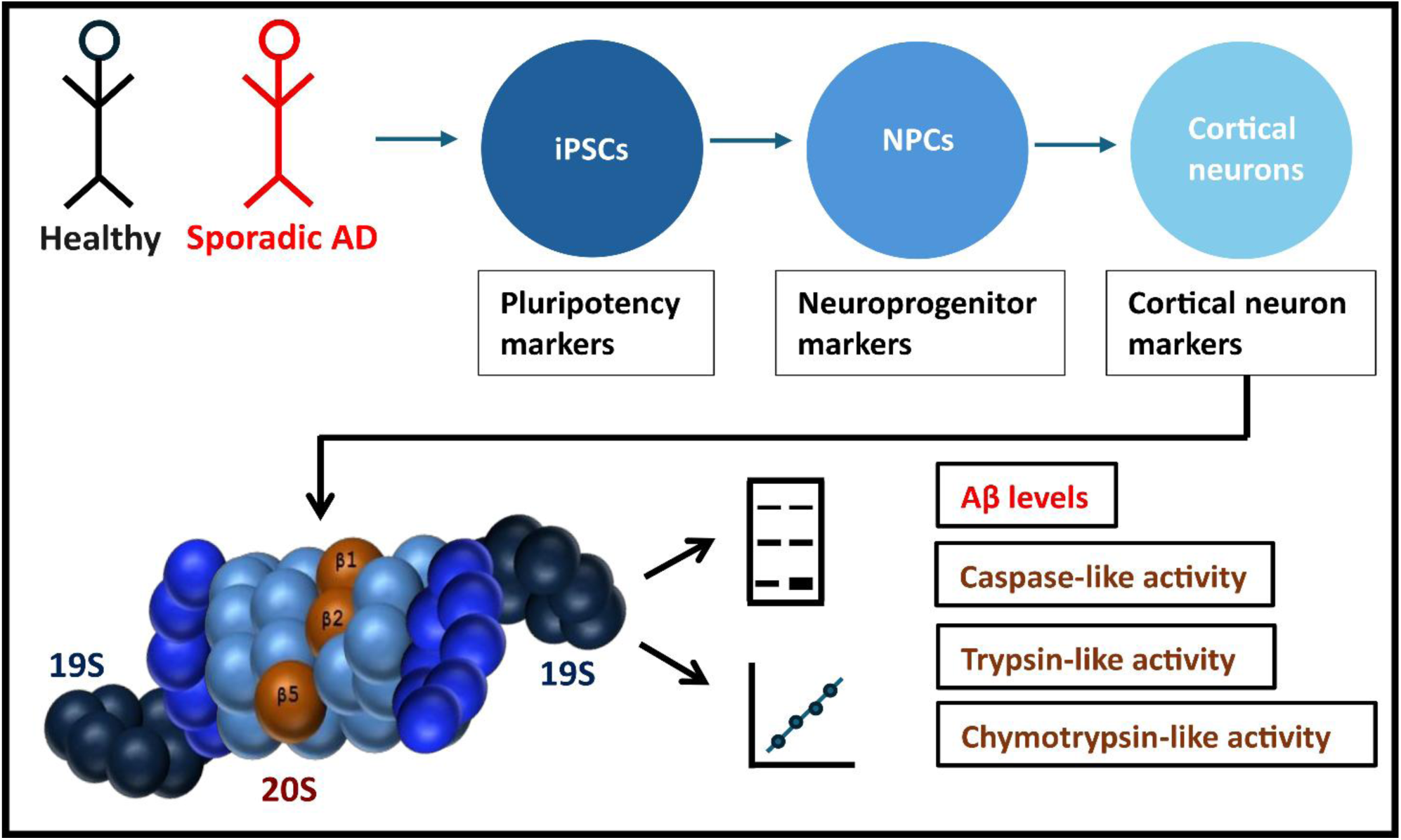

## 1. Introduction

Alzheimer’s disease (AD) is a major cause of disability and death especially in the elderly and currently has no cure [1]. Symptomatically, AD is characterized by progressive cognitive decline and behavioral changes that deteriorate quality of life. Neuropathologically, AD is characterized by the extracellular deposition of amyloid-β (Aβ) peptide in the form of diffuse and neuritic plaques and the accumulation of intraneuronal neurofibrillary tangles consisting of aggregated hyperphosphorylated tau protein [2–3]. Brain Aβ burden has been regarded as a critical player in the disease etiology with a protracted preclinical phase of up to 20 years that is marked by deposition of Aβ preceding the onset of clinical symptoms [4]. Sporadic AD (sAD) accounts for more than 90% of all AD cases but understanding its etiology and pathogenesis continues to be challenging. These challenges arise from limited access and availability of viable neuronal cells from sAD patients due to ethical and practical reasons. Also, the reliance on transgenic animal models, while valuable for mechanistic insights, has proven inadequate for recapitulating human sAD pathology especially at the cellular level and for translating discoveries into effective human therapies [5].

Human induced pluripotent stem cell (iPSC) technology has enabled the generation of translational cellular models for better understanding of the molecular underpinnings of sAD. Human iPSC-derived neuronal models offer advantages over conventional transgenic animal models. These advantages include preservation of human-specific cellular and molecular features and the great potential for personalized medicine [6]. A major advantage is the ability to generate unlimited numbers of cortical neurons from patient iPSCs for accurate analyses of native enzymatic activities which is challenging with post-mortem brain samples. Human iPSC-derived neurons have been shown to recapitulate key cellular phenotypes of AD including increased Aβ production, tau hyperphosphorylation, and cellular dysfunction [6–8]. These cellular models display perturbations in multiple cellular pathways, including the ubiquitin-proteasome system (UPS) [9–10]. The proteasome is the major multi-subunit and multi-catalytic protein complex of the UPS. It is vital for maintaining intracellular protein quality control by removing misfolded or aggregate-prone damaged proteins in eukaryotic cells [11]. It is composed of the barrel-shaped 20S core particle (CP) and the 19S regulatory particle (RP). The 20S CP contains the 3 proteolytic active sites (β1, β2, and β5), generally classified as caspase-like for β1, trypsin-like for β2, and chymotrypsin-like for β5. Collectively, these proteolytic sites enable the degradation of a wide variety of substrates following covalent tagging with polyubiquitin chains [11–12]. The 19S regulatory particle (RP) recognizes the ubiquitin-tagged substrates, cleaves the ubiquitin chains, unfolds the substrates, and translocates them into the proteolytic chamber of the 20S CP. The 20S CP (∼700 kDa) and the 19S RP (∼900 kDa) interact dynamically to form different proteasome assemblies.

Association of the 19S RP with either one or both ends of the 20S CP results in the formation of the 26S (RP1-CP) or 30S (RP2-CP) proteasomes, respectively. The RP2-CP, RP1-CP, and CP proteasomes, as well as various assembly intermediates, all coexist in the cell [11–12]. Due to its essential proteostatic role, the proteasome is critical for the normal functioning of long-lived post-mitotic cells like the neurons that cannot regenerate.

Increasing evidence suggests that proteasome dysfunction plays a central role in the pathogenesis of AD [13–15]. Although proteasome activity has been investigated in various AD models, these models have mainly represented late-stage conditions, leaving a void in the understanding of proteasome assemblies, activity, and dynamics in early-stage conditions. The present study aimed to characterize the changes in proteasome assemblies, proteolytic activities, and possible compensatory adaptations occurring in early-stage sAD.

## 2. Materials and Methods

### 2.1 Human induced pluripotent stem cells

The human induced pluripotent stem cell (iPSCs) used in this project were provided by the University of California Alzheimer’s Disease Research Center (UCI-ADRC) and the Institute for Memory Impairments and Neurological Disorders. All iPSC work was performed with the approval of the Institution Review Board (IRB) of the Helsinki Institute of Life Sciences at the University of Helsinki (Finland). Human iPSCs were obtained from the fibroblasts of 3 normal subjects and 3 patients diagnosed with sporadic Alzheimer’s disease. All the iPSC lines used in this study are listed in Table S1 and cultured as described in the supplementary method.

### 2.2 Neural induction of iPSCs

The iPSCs were induced into neural progenitor cells (NPCs) followed by terminal differentiation into mature neurons using a well-established dual SMAD inhibition protocol with modifications [16]. Briefly, iPSCs were plated at high density and neural induction was initiated upon reaching 80-90% confluence on Matrigel-coated dishes by addition of neural induction medium (NIM). The NIM consists of 1:1 DMEM/F12 and neurobasal medium, 1x N-2 supplement, 1x B-27 supplement minus vitamin A, 1x Sodium Pyruvate, 1x nonessential amino acids [NEAA], 1x GlutaMAX, 100 μM β-mercaptoethanol, 5 μg/ml insulin (Thermo Fisher Scientific), and 0.2 μM LDN193189, 10 μM SB431542, and 5μM XAV939 (MedChemExpress) to block the BMP/TGF-β and WNT signaling pathways. Medium was changed daily until neural rosettes appeared.

### 2.3 Neuronal differentiation

Neural progenitor cells were isolated from neural rosettes and expanded in NPC maintenance medium composed of 1:1 DMEM/F12 and neurobasal medium, 1x N-2 supplement, 1x B-27 supplement minus vitamin A, 1x Sodium Pyruvate, 1x NEAA, 1x GlutaMAX, 100 μM β-mercaptoethanol, 5 μg/ml insulin, 20 ng/ml bFGF, and 20ng/ml EGF (Thermo Fisher Scientific). For terminal neuronal differentiation, NPCs were dissociated and singularized on poly-L-ornithine, laminin, and fibronectin triple-coated coverslips or culture plates in neuronal differentiation medium composed of 1:1 vol/vol of DMEM/F12 and neurobasal medium, 1x N-2 supplement, 1x B-27 supplement minus vitamin A, 1x Sodium Pyruvate, 1x NEAA, 1x GlutaMAX, 100 μM β-Mercaptoethanol, 10μg/ml insulin, 200μM ascorbic acid, 100μM dibutyryl cAMP, 20ng/ml BDNF, 20ng/ml NT-3, and 20ng/ml GDNF to promote the phenotypic specification and maturation into cortical neurons.

Proliferating cells were eliminated by adding 3 μM Cytarabine to the differentiation medium on DIV 2 of neuronal differentiation for 48 hours. Cells were plated at 300,000/well and 10,000/well in 24-well plates for high density (protein analysis) and low density (microscopy) cultures respectively. Half-medium change was performed every 2 days for the first 2 weeks. Thereafter, half-medium change was made every 3-4 days, and neurons were maintained for about 8 weeks.

### 2.4 Immunofluorescence Staining

Cells were fixed with 4% paraformaldehyde in phosphate-buffered saline (PBS) for 20 minutes at room temperature, followed by permeabilization with 0.1% Triton X-100 in PBS for 10 minutes. Non-specific binding was blocked using 5% normal goat serum and 1% bovine serum albumin in PBS for 1 hour at room temperature. Pluripotency was validated by established markers including OCT4, NANOG, and SSEA4. The expressions of OCT4, NANOG, and SSEA4 were used to confirm the undifferentiated state and an indication of pluripotency of the human control and AD iPSCs lines. Pluripotency markers OCT4 and NANOG are transcription factors robustly expressed in the nuclei of undifferentiated stem cells and play pivotal roles in self-renewal and pluripotency. The glycolipid antigen SSEA4 is expressed on the surfaces of human embryonic and induced pluripotent stem cells and is also used as a marker of pluripotency. Conversion of control and sAD human iPSCs into dorsal NPCs following dual SMAD inhibition was verified by the expression of key markers NESTIN and PAX6 to monitor for successful neural induction and maintenance of progenitor identity.

Neuronal identity was confirmed by markers βIII-tubulin (TUJ1) and MAP2, while cortical identity was validated using TBR1 as cortical layer marker. Primary antibodies were applied overnight at 4°C in blocking solution at the following concentrations: OCT4 (1:500, DSHB), NANOG (1:400, DSHB), SSEA4 (1:200, Abcam), PAX6 (1:300, Proteintech), NESTIN (1:500, Proteintech), TUJ1 (1:1000, Biolegend), MAP2 (1:800, Aves Lab), and TBR1(1:500, Proteintech). Following primary antibody incubation, cells were washed 3x with PBS and incubated with appropriate Alexa Fluor-conjugated secondary antibodies (1:1000, ThermoFisher) for 1 hour at room temperature in the dark. Nuclei were counterstained with DAPI (1 μg/ml) for 10 minutes. Coverslips were mounted using Fluoromount-G antifade (SouthernBiotech) and fluorescent images were obtained using an EVOS M5000 microscope.

### 2.5 Protein extraction and western blot analysis

Total protein extracts were prepared from iPSC-derived cortical neurons using RIPA buffer (50 mM Tris-HCl pH 7.4, 150 mM NaCl, 1% NP-40, 0.5% sodium deoxycholate, 1% SDS) supplemented with protease and phosphatase inhibitor cocktail (ThermoFisher). Cells were lysed on ice for 30 minutes with periodic vortexing, followed by centrifugation at 10,000xg for 10 minutes at 4°C to remove debris. Total protein concentrations were determined using the BCA protein assay kit (ThermoFisher). For western blot analysis, 15 μg of protein were separated by SDS-PAGE using 4-12% Bis-tris gradient gel systems (GeneScript) and transferred to PVDF membranes (BioRad) using a semi-dry transfer system. Membranes were blocked with 3% non-fat dry milk in phosphate-buffered saline (PBS) with 0.1% Tween-20 (PBST) for 1 hour at room temperature, followed by overnight incubation at 4°C with primary antibodies against Proteasome subunit beta type-5 (PSMB5, 1:1000, Proteintech) and GAPDH (1:5000, Proteintech) as a loading control. After washing with PBST, membranes were incubated with horseradish peroxidase-conjugated anti-mouse and anti-rabbit IgG secondary antibodies (1:10000, Proteintech) for 2 hours at room temperature. Protein bands were visualized using enhanced chemiluminescence (ECL) detection reagent (Millipore) and imaged using a G-Box imaging system (Syngene).

Densitometric analysis was performed using ImageJ software, and PSMB5 levels were normalized to GAPDH loading controls.

### 2.6 Enzyme-Linked Immunosorbent Assay (ELISA)

Intraneuronal levels of total amyloid-β peptides (Aβ) were quantified via an in-house sandwich ELISA protocol using Aβ1-17 as standard, anti-APP/Aβ monoclonal antibody clone 6E10 (Biolegend) as the capture antibody and anti-Aβ monoclonal antibody clone 82E1 (biotinylated) (Immuno-Biological Laboratories) as the detection antibody. The clone 6E10 immunogen spans amino acids 1–16 of Aβ, so it recognizes Aβ peptides, full-length APP and APP fragments because the Aβ1-16 sequence is embedded within the APP ectodomain [17]. The clone 82E1 specifically requires a free N-terminus generated via β-secretase cleavage of APP. Due to its strict binding requirement for an unmodified Asp1, 82E1 is highly specific for detecting bona fide Aβ peptides generated by sequential β- and γ-secretase cleavage of APP. Therefore, clone 82E1 does not recognize full-length APP, or any N-terminal Aβ species beginning at positions 2 or 3. Importantly, clone 82E1 allows unambiguous detection of total Aβ fragments in complex biological samples, and reacts with soluble and fibrillar Aβ to a similar degree [18]. Due to the broad cross-reactivities of clone 6E10 with full-length APP, APP fragments, and Aβ, it was used as the capture antibody in this study. The clone 82E1, owing to its specificity for Aβ, was used as the detection antibody. Human iPSC-derived cortical neurons were prepared and lysed in RIPA buffer. An ELISA plate was coated with 6E10 diluted in 50 mM NaHCO_3_ pH 9.6 to a concentration of 1µg/ml and incubated for 72 hours at 4°C with gentle shaking. Blocking was done using 1% bovine serum albumin (BSA) dissolved in PBST. Standards and samples were added in duplicates and incubated for 48h at 4°C with gentle shaking. The detection antibody, biotinylated 82E1, diluted in 0.1% BSA-PBST to a concentration of 1µg/ml was added to the plate and incubated for 24h at 4 °C with gentle shaking. Streptavidin-HRP conjugate diluted 1:10,000 in 0.1% BSA-PBST was added and incubated for another 24h at 4 °C with gentle shaking. Color development was achieved by adding TMB substrate and incubated at 37°C for 10min. The reaction was quenched by 1M HCl and absorbance was measured at 450 nm using a Varioskan Flash microplate (ThermoFisher).

### 2.8 Native PAGE analysis of proteasome assemblies

The composition of active proteasomes was analyzed using the fluorogenic peptide substrate Suc-LLVY-AMC using native in-gel activity assays. This approach enables the separation and visualization of the different proteasome complexes without disrupting their native structures and catalytic functions [19]. Neurons were lysed in native lysis buffer (50 mM Tris-HCl pH 7.5, 150 mM NaCl, 5 mM MgCl_2_, 0.5% NP-40, 2 mM ATP, 1 mM DTT) supplemented with protease and phosphatase inhibitor cocktail. Lysates were incubated on ice for 30 minutes and centrifuged at 10,000xg for 10 minutes. Total protein concentration was determined using the BCA (ThermoFisher). Native PAGE was performed using a 4.5% Tris-acetate gel with a 3% Tris stacking gel. Protein samples (30μg) were mixed with native sample buffer and loaded onto the gel. Purified human 20S was loaded as a standard. Electrophoresis was conducted at 4°C with constant voltage (150V) for 5.5 hours to allow adequate separation of proteasome complexes. For in-gel proteasome activity assay, gels were incubated in proteasome activity buffer (50 mM Tris-HCl pH 7.5, 5 mM MgCl_2_, 1 mM ATP, 0.5mM DTT) containing 24μM Suc-LLVY-AMC ( Peptide Institute) for chymotrypsin-like activity detection in the absence and presence of 0.035% SDS. Gels were incubated at 37°C for 30 minutes, and fluorescent bands corresponding to active proteasome complexes were visualized using the G-Box imaging system (Syngene). The positions of 30S (doubly-capped), 26S (singly-capped), and 20S (uncapped) proteasome complexes were identified based on their characteristic migration patterns and molecular weights. Densitometric analysis was performed to quantify the relative abundance of the active proteasome complexes as a percentage of the control by setting the value obtained in the control at 100%.

### 2.9 Proteasome activity-based profiling

Activity-based profiling (ABP) was used to label the 3 catalytic β-subunits (β1, β2, and β5) of the proteasome in a single step, providing a global snapshot of proteasome activity as well as global allostery among the β-subunits (see supplementary methods for details).

### 2.10 In-solution kinetic activity assays

For enhanced precision, the behaviors of the 3 proteolytic sites of the proteasomes were individually investigated using fluorogenic substrates specific for each of the β1, β2, and β5 catalytic subunits as described in the supplementary method.

### 2.11 Statistical Analysis

Analyses were performed using Microsoft 365 Excel and KyPlot 6.0. Data are presented as mean ± standard error of the mean (SEM) unless otherwise indicated. Statistical comparisons between groups were performed using Welch’s t-tests (two-tail). Statistical significance was set at p < 0.05 (* p < 0.05, ** p < 0.01, and *** p < 0.001).

## 3. Results

### 3.1 Control and sporadic AD human iPSCs robustly express markers of pluripotency

Human iPSCs derived from normal and sAD subjects (Table S1) were subjected to the dual-SMAD neuronal differentiation protocol (Figure 1A). Phase-contrast microscopy images revealed that iPSCs from both control and sAD lines had the typical morphology of pluripotent stem cells. The iPSCs propagated well in adherent monolayer cultures with round shape and uniform morphology. They were clustered together in colonies and tightly arranged without obvious cell boundaries. Colonies were island-like with clear edges and appeared uniform and identical. High nuclear-to-cytoplasmic ratio was observed with larger nuclei and relatively less cytoplasm (Fig. 1B). Immunofluorescence revealed robust expressions of OCT4, NANOG, and SSEA4 in both control and sAD iPSCs. Immunostaining revealed strong nuclear staining for both OCT4 and NANOG (Figure 1C), with SSEA4 displaying plasma membrane staining (Figure S1A). Double immunofluorescence analysis established that all nuclei were positive for both OCT4 and NANOG. The morphology and staining intensity of the pluripotency markers were comparable between control and sAD lines, indicating successful reprogramming and maintenance of pluripotent state. The morphology of all iPSCs from control and sAD lines was consistent across multiple passages, ensuring the stability of the starting cell populations.

**Figure 1:**
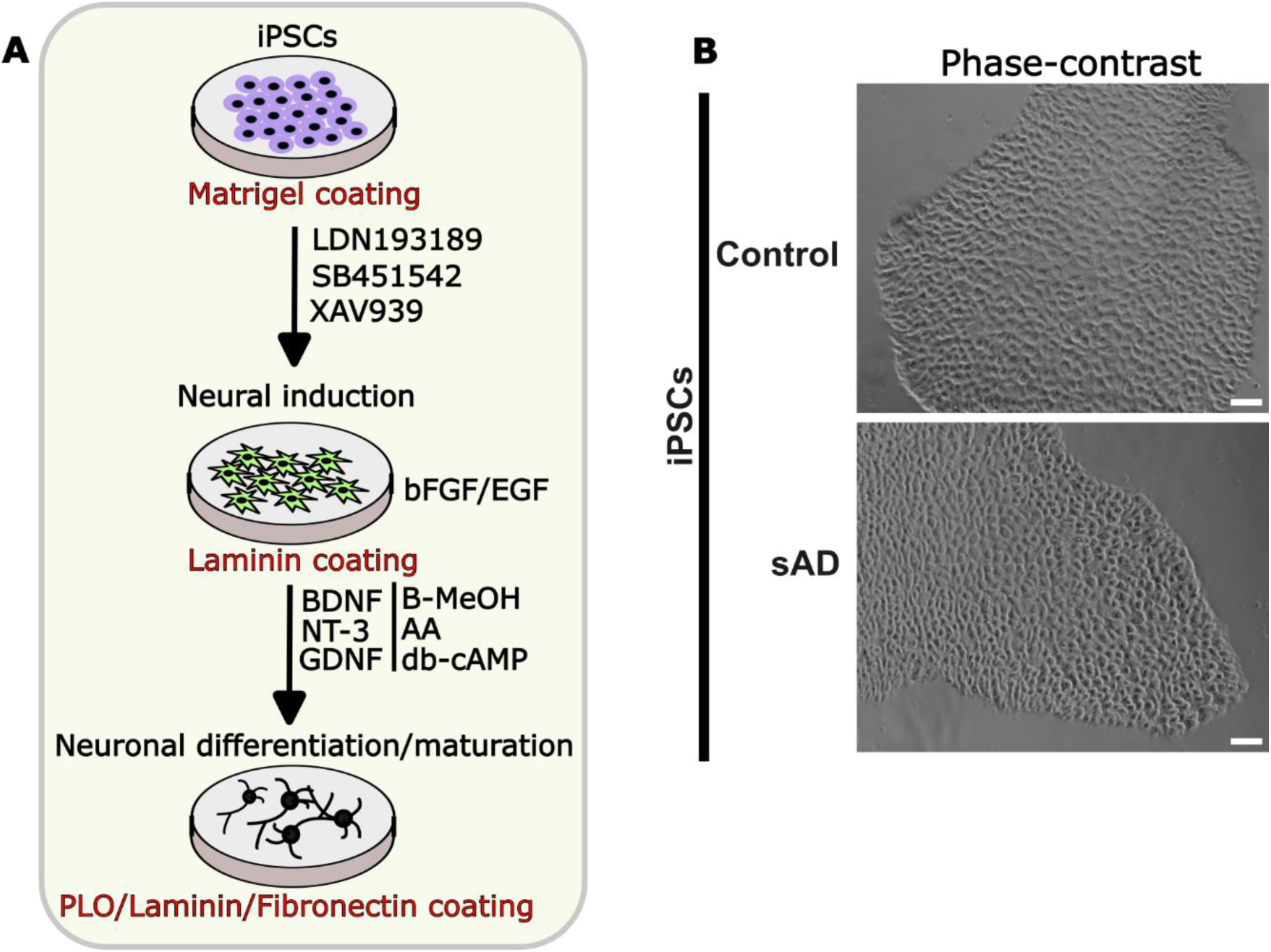
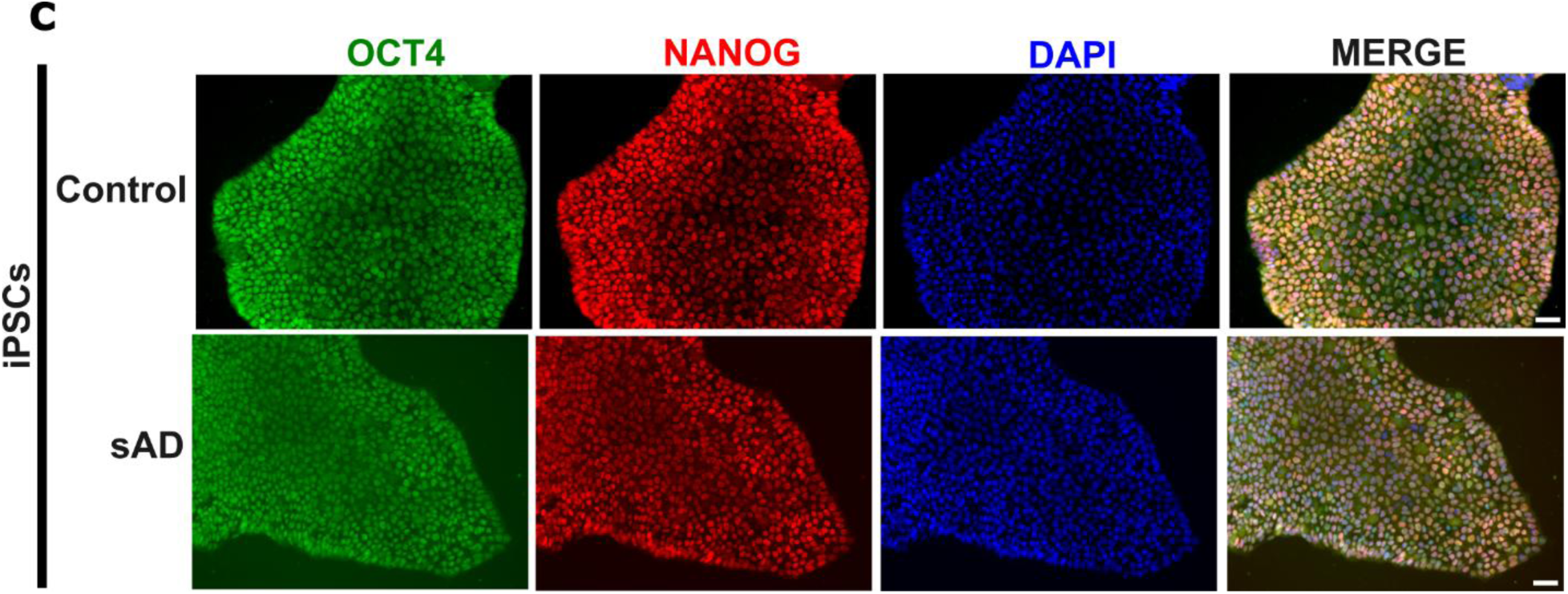
Control and AD iPSCs expresses markers of pluripotency. (A) Schematic overview of the procedures for generating cortical neurons from human induced pluripotent stem cells (iPSCs) using the dual-SMAD protocol. (B) Representative phase-contrast microscopy of human control and sAD iPSCs showing typical colony morphology. Scale bar = 50µm. (C) Representative IHC of control and sAD iPSCs expressing pluripotency markers OCT4 (green) and NANOG (red). Nuclei were stained by DAPI. Scale bar = 20µm.

### 3.2 Dual SMAD inhibition promotes induction of control and sporadic AD iPSCs into dorsal NPCs and forebrain cortical neurons

Immunofluorescence staining of the dorsal NPCs showed strong expression of NESTIN in the cytoplasm (Figure 2A), reflecting their neuroectodermal and multipotent identities [20]. A high number of NESTIN-positive cells also displayed robust expression of the transcription factor PAX6 in their nuclei (Figure 2A), confirming their dorsal identity. Immunostaining further revealed the formation of characteristic neural rosettes which are elongated polarized structures containing radially organized columnar cells [21] in NPC cultures derived from both control and sAD iPSC lines. These rosettes, which strongly co-express NESTIN and PAX6, have been shown to correspond to the neuroepithelial stem cells of the developing cortex based on their strong apico-basal epithelial polarity [22]. The co-expression of NESTIN and PAX6 in the dorsal NPCs indicates efficient neural induction (Figure 2A). This also confirms that sAD has no observable effect on neural induction of iPSCs into NPCs following dual SMAD inhibition, in agreement with a previous study [8].

**Figure 2:**
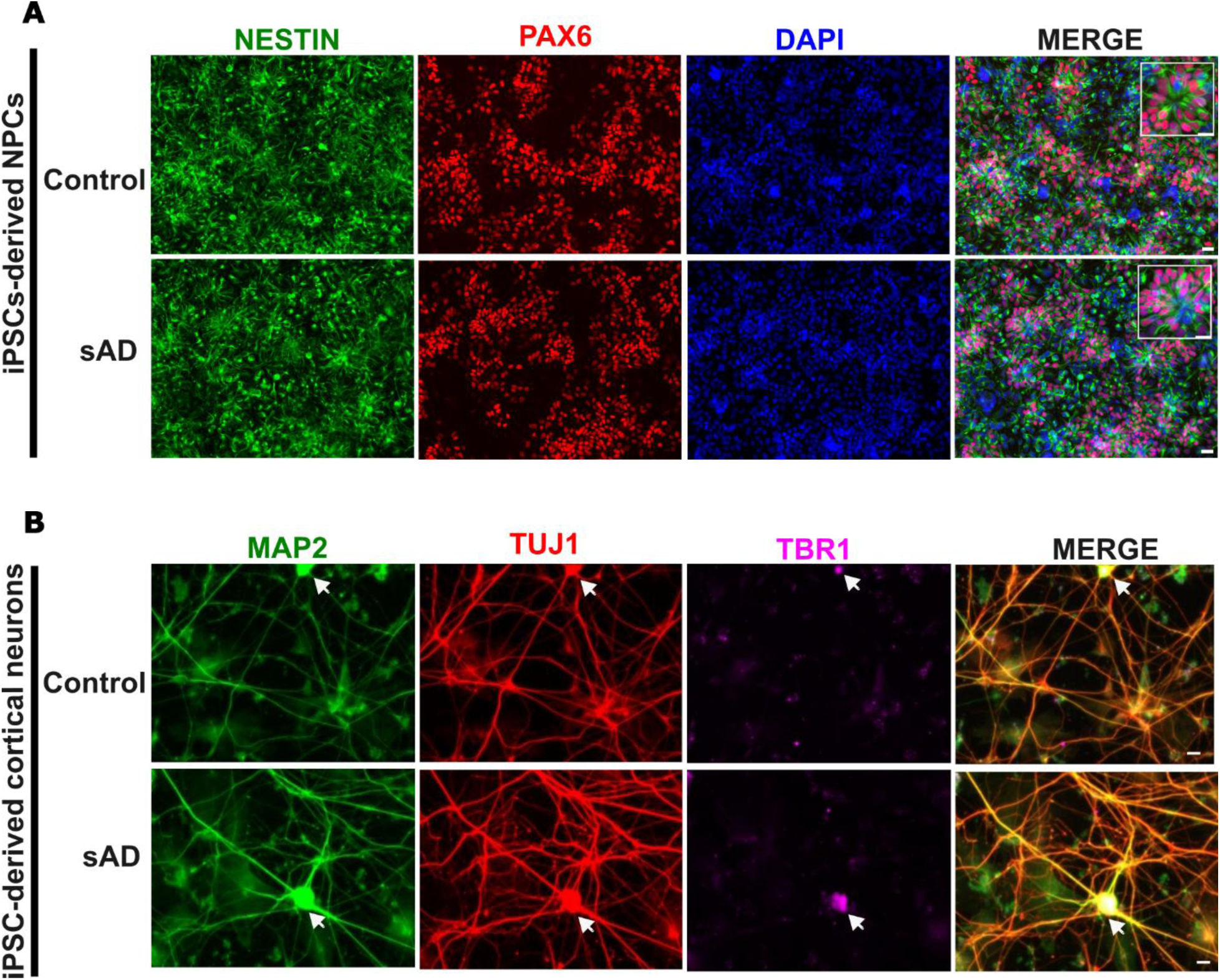
Immunofluorescence characterizations of iPSC-derived neuroprogenitor cells (NPCs) and cortical neurons. (A) Representative IHC of control and sAD neural stem cells expressing neuroprogenitor markers NESTIN (green) and PAX6 (red). Nuclei were stained by DAPI. The insets show the radially organized and polarized structures characteristic neural rosettes. Scale bar = 20µm. (B) Representative IHC of control and sAD iPSC-derived cortical neurons expressing mature neuronal makers MAP2 (green) and TUJ1 (red), as well as cortical layer marker TBR1 (magenta). Scale bar = 20µm.

The dorsal NPCs from both control and sAD lines acted as functional progenitors by differentiating into homogenous population of mature cortical neurons as evidenced by extensive neurite outgrowth and expression of neuronal markers. Immunofluorescence staining revealed strong expressions of TUBB3 (TUJ1) and MAP2 in neurons derived from both control and sAD dorsal NPCs (Figure 2B). These neurons displayed strong cytoplasmic immunoreactivity of the cytoskeletal protein TUBB3 by the TUJ1 antibody, staining the soma and the fine details of the axon and dendrites. Another cytoskeletal protein MAP2, a more mature marker, was strongly immunoreactive in the soma and dendrites of neurons. To characterize the cortical identity of the neurons differentiated from the dorsal NPCs, the deep cortical layer (V and VI) marker TBR1 was used (Figure 2B). The expression of these key markers showed no observable differences between the control and sAD lines, indicating that the disease background has no observable effect on the terminal differentiation of iPSCs into cortical neurons using the experimental conditions described in this study.

### 3.3 Total intraneuronal Aβ level was elevated in sporadic AD iPSC-derived cortical neurons

The ELISA analysis revealed a significant increase (p = 0.0124) in the intraneuronal level of total Aβ in the sAD iPSC-derived cortical neurons compared to the control (Figure 3). The mean value of 110pg/ml total intraneuronal Aβ level obtained for the sAD neurons falls within the previously reported range of 50-1800 pg/ml [23]. Aβ secretion from iPSC-derived neurons has been reported to be highly maturation-dependent, increasing depending on the duration of culture [8]. Thus, the total intraneuronal Aβ concentration obtained in this study suggests that these, under our experimental conditions, may be representative of early-stage sAD neuropathology. Quality control metrics revealed an overall intra-assay % coefficient of variation (%CV) of 2.7%. Typically, an intra-assay %CV of 10% or less is considered satisfactory, reflecting the reliability of the ELISA [24]. An intra-assay %CV of < 10% demonstrates high assay precision and reproducibility for total intraneuronal Aβ measurements. This result demonstrates that these sAD iPSC-derived cortical neurons recapitulate a key sAD feature involving the elevation of intraneuronal Aβ levels [17].

**Figure 3:**
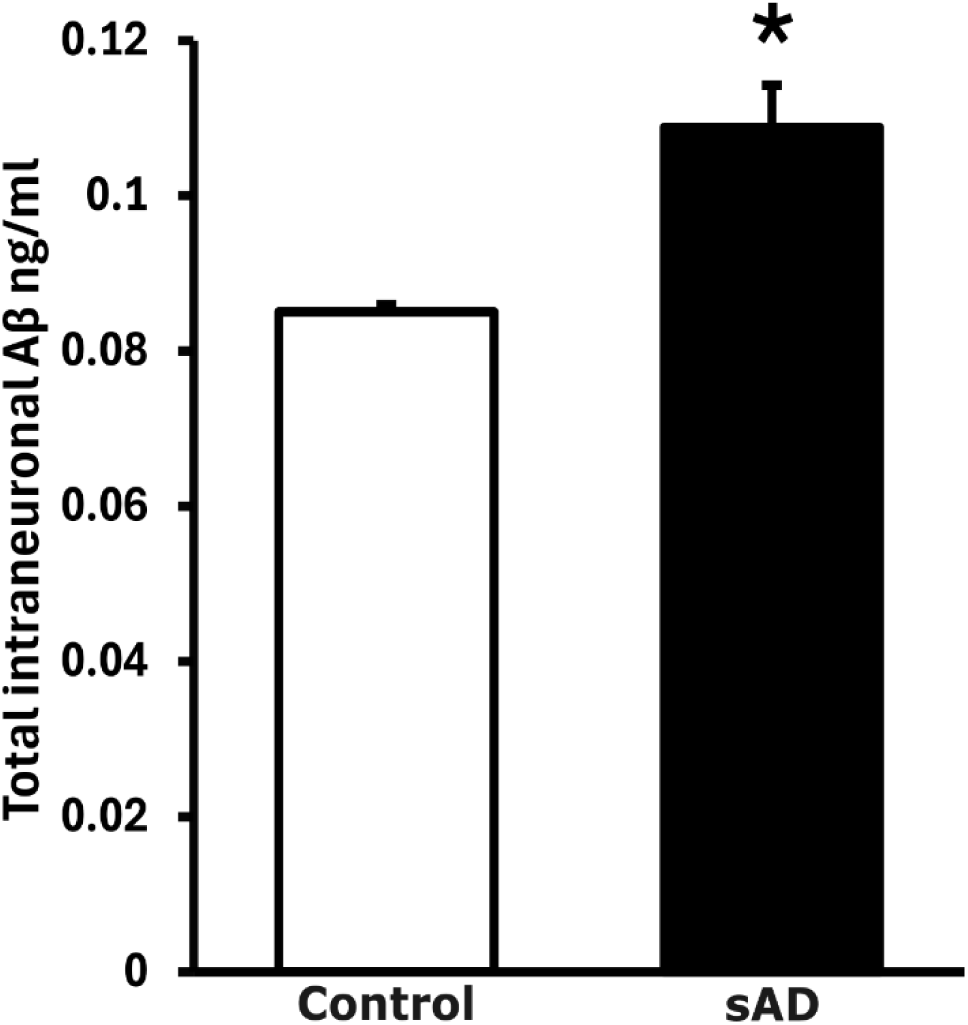
ELISA quantification of the of total intraneuronal Aβ in control and sAD cortical neurons. Measurement of total intraneuronal Aβ in control and sAD iPSC-derived cortical neurons after 8 weeks in culture showed increased levels in sAD. Data represents mean ± SEM (n = 3), p = 0.0124. T test was performed to evaluate significance between control and sAD cortical neurons at the same point of differentiation (*p < 0.05).

### 3.4 Active 20S proteasome assemblies are increased in sporadic AD iPSC neurons

Analysis of the active proteasome assemblies showed that lysates from both control and sAD neurons exhibited proteasome chymotrypsin-like (CT-L) activity at bands corresponding to double-capped 30S proteasome (∼2.9 MDa), single-capped 26S proteasome (∼2 MDa), and uncapped 20S proteasome (∼700 kDa) (Fig. 4A-B). The 30S, 26S and 20S proteasome complexes were all present in both the control and AD neurons (Fig. 4A-B). There were no significant differences in the levels of active 30S (p = 0.7355) and 26S (p = 0.7982) complexes in the control and sAD neurons (Fig. 4A, C, D). The 30S and 26S proteasomes play central roles in degrading substrates in ATP-dependent manner [25], thus suggesting that the substrate recognition function of the 26S/30S proteasomes might not be impaired in early-stage sAD. In contrast, the sAD neurons showed an increased CT-L activity at the 20S band meaning a significantly higher level of active 20S proteasomes compared to the control (p = 0.0177, Fig. 4B, E). The elevation in 20S proteasomes activity in sAD neurons has important functional implications because they can degrade substrates in ATP-independent manner compared with the 26S/30S proteasomes [26].

**Figure 4:**
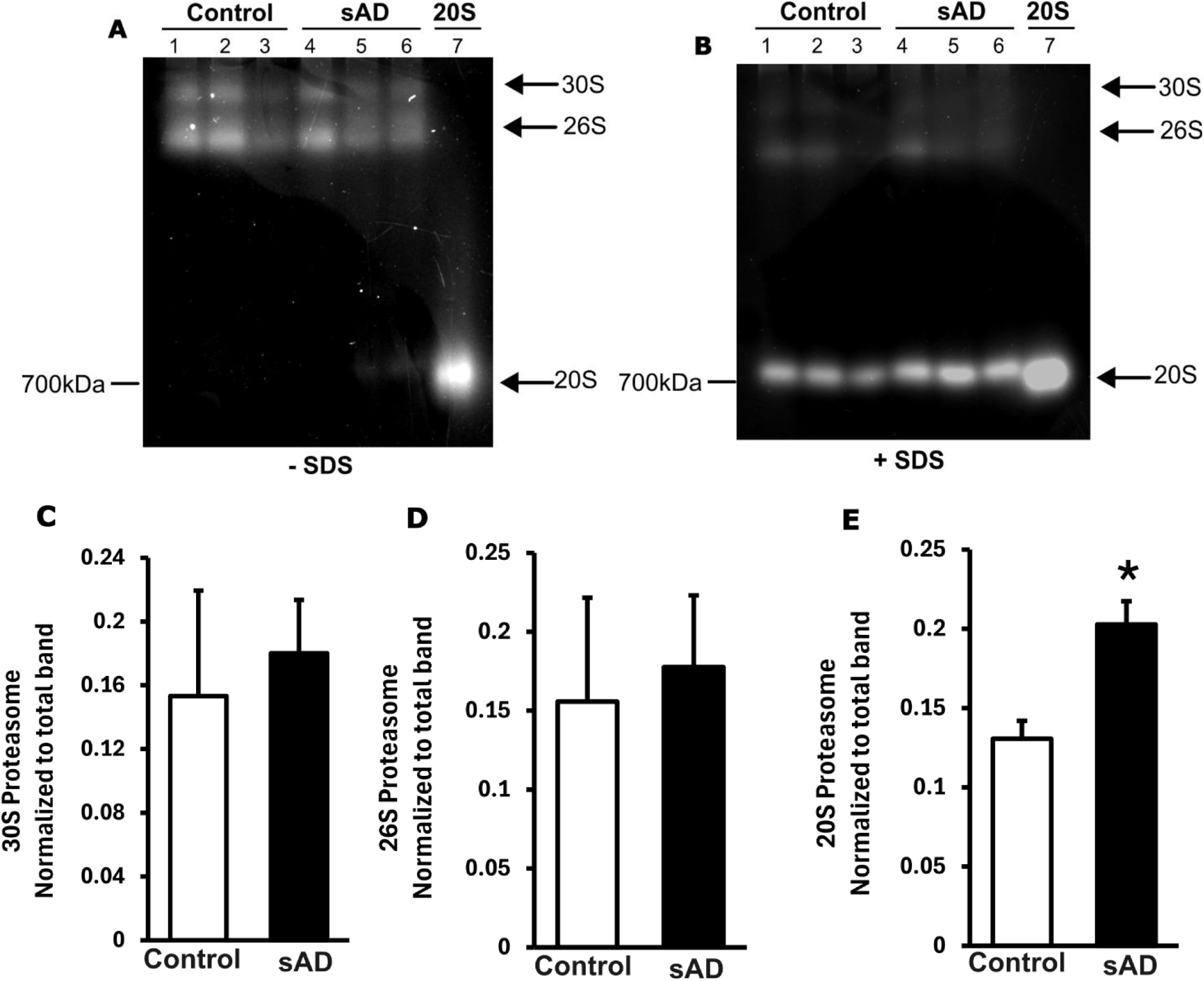
Composition of active proteasome subtypes in control and sAD cortical neurons. (A), (B) Native activity gels of control and sAD neurons in the absence/presence of SDS revealed bands corresponding to 30S, 26S, and 20S proteasome assemblies. Purified 20S proteasome with molecular weight of 700kDa was used as marker. (C-D) The levels of active 30S and 26S revealed no significant changes between control and sAD neurons, p = 0.7355 and p = 0.7982 respectively. Data represents mean ± SEM (n = 3). (E) The sAD neurons have higher composition of 20S proteasome sub-type compared to control, p=0.0177. Data represents mean ± SEM (n = 3).

Thus, the increased level of active 20S proteasomes in the sAD neurons may suggest a compensatory response to increased proteotoxic stress at the early stage of the disease, in agreement with earlier studies showing increased 20S proteasome in response to various cellular stressors [26–28].

### 3.5 Activity-based profiling reveals specific 20S proteasome subunit elevation in sporadic AD neurons

The activity-based labelling of control and sAD cortical neuron lysates revealed distinct fluorescent band labeling by the Me4BodipyFL-Ahx3Leu3VS probe (Fig. 5A) corresponding to the β1, β2, and β5 subunits of the 20S proteasome (Fig. 5B). All 3 catalytic subunits activities in both control and sAD neurons displayed the typical band patterns consistent with previous studies [29–30]. The β5 subunit activity demonstrated the most intense labelling compared to β1 and β2 in both control and sAD neurons (Fig. 5B), suggesting an important role for the CT-L activity of the 20S proteasome in response to proteotoxic stress [31–32]. This also supports earlier reports that revealed the β5 subunit activity as the initial and rate-limiting step in overall substrate degradation by the proteasome [33]. Analysis revealed significant increases in the β1 (p = 0.004), β2 (p = 0.023), and β5 (p = 0.029) subunit activities in the sAD neurons compared to control (Fig. 5C-E). These results indicate that proteasome activation in the sAD neurons occurs similarly across all catalytic activities, pointing a concerted compensatory response to proteotoxic stress probably owing to intraneuronal Aβ elevation. To reinforce these findings, immunoblotting was performed to specifically quantify the protein abundance of PSMB5 which is responsible for the β5 subunit activity of the proteasome. Between control and sAD neurons, the total protein levels of PSMB5 did not reveal any significant difference (p = 0.431, Fig. 5F). This indicates that the elevated activity-based labeling of the β5 subunit reflects an increase in catalytic activity rather than increased subunit expression.

**Figure 5:**
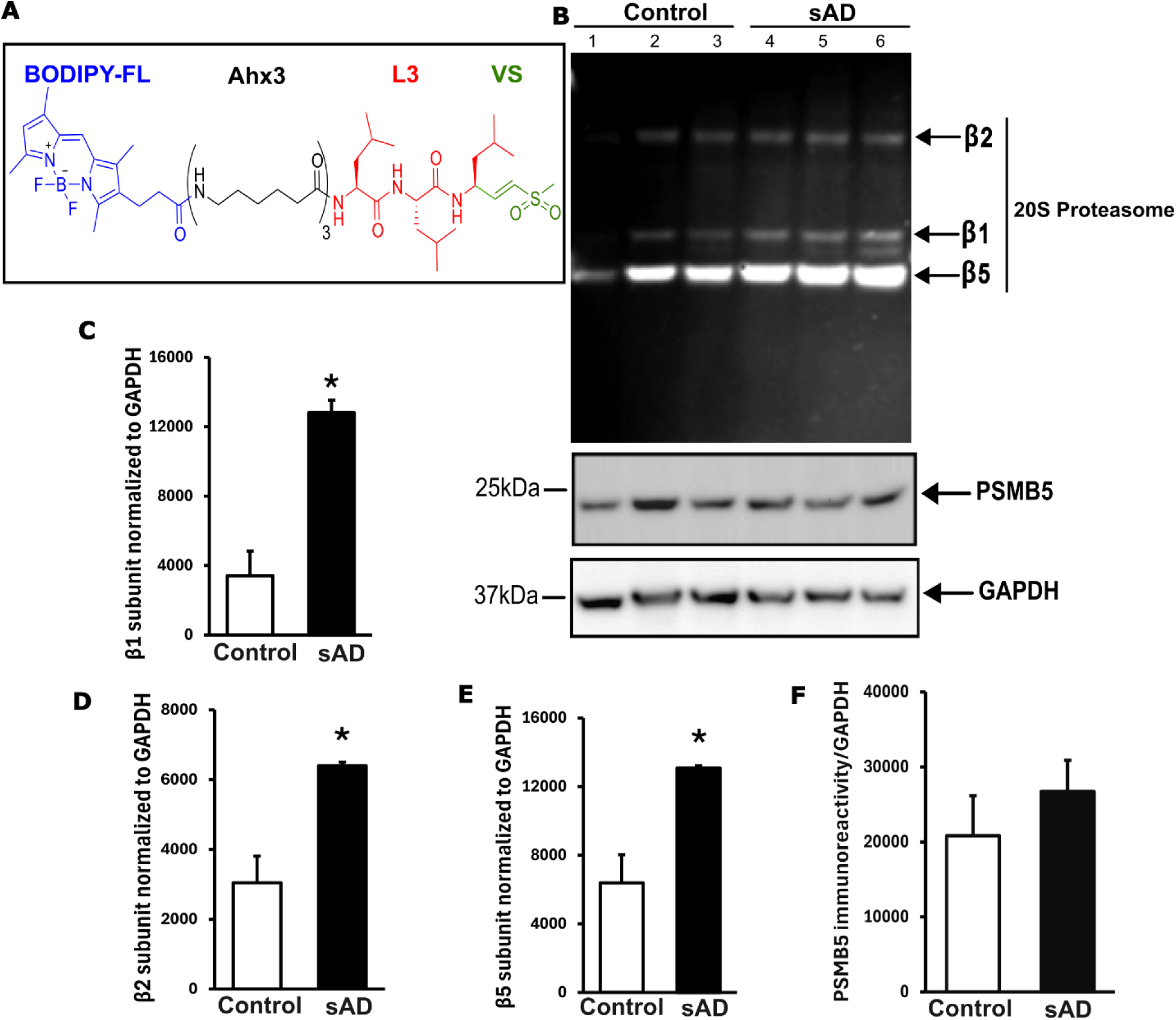
Activity-based profiling of β1, β2, and β5 proteolytic sites in control and sAD cortical neurons. (A) Me4BodipyFL-Ahx3Leu3VS probe with the fluorescent BODIPY-FL and reactive vinyl sulfone (VS) moieties. (B) Migration patterns of the β1, β2, and β5 sites of active proteasomes in control and cortical sAD neurons (upper gel). PSMB5 and GAPDH quantification by western blot (middle and lower gels). (C-E) Higher activity of β1, β2, and β5 sites in sAD cortical neurons compared to control, p = 0.004, p = 0.023, and p = 0.029 respectively. Data represents mean ± SEM (n = 3). (F) PSMB5 protein expressions in control and sAD cortical neurons revealed no significant difference, p = 0.431. Data represents mean ± SEM (n = 3).

### 3.6 Kinetic analysis validates increased proteolytic activities of the proteasome in sporadic AD neurons

The significant elevation of the proteolytic activities of the β1, β2, and β5 subunits was further supported by individual profiling of these catalytic subunits using fluorogenic peptides specific for the CT-L (chymotrypsin-like) activity (β5), T-L (trypsin-like) activity (β2), and C-L (caspase-like) activity (β1) in kinetic assays. Slope analysis of the β1 C-L proteolytic activity progress curves, examined by Z-LLE-AMC, revealed a significant increase of 100% (p = 0.006) in sAD neurons compared to control (Fig. S6A-C). The slope of β2 T-L proteolytic activity progress curves, examined by Boc-LRR-AMC, showed a significant increase of 144.83% (p = 0.021) in sAD neurons compared to control (Fig. S6B, D). The slope of β5 CT-L proteolytic activity progress curves resulted in a significant increase of 55.75% (p = 0.037) in sAD neurons compared to control (Fig. S6B, E). The AUC analysis of the β1, β2, and β5 progress curves all revealed significant increases of 68.86% (p = 0.007), 61.01% (p = 0.018), and 113.90% (p = 0.007) respectively in the sAD neurons compared to the controls (Fig. S6F-H). These results are consistent with the activity-based profiling results. Together, these results suggest that the slope and AUC measure proteolytic activities of the proteasome differently but complementarily, providing added robustness that neither alone can achieve.

## 4. Discussion

This study investigated the proteasomal proteolytic activity in a cellular model of sAD using human iPSC-derived cortical neurons as cellular models of the early stage of the disease. Earlier studies using transgenic animals and post-mortem clinical samples have reported the inhibition of the proteasome activity due to aggregates of Aβ and Tau in AD [9–10].

However, the models used in virtually all these studies represent late-stage AD pathology due to the forced aggressive Aβ accumulation in transgenic models and the long pre-symptomatic Aβ accumulation phase in patients. Thus, the status of proteasomal activity in early-stage sAD is currently unexplored. Here, we have utilized orthogonal tools to investigate the compositions and activity of the proteasome complexes in cortical neurons differentiated from control and sAD patients iPSCs.

The iPSC-derived cortical neurons generated in this study passed through stages reminiscent of in vivo neurogenesis and displayed key developmental markers of pluripotency (OCT4, NANOG, and SSEA4), neuroprogenitor (NES and PAX6), and cortical identity (TUJ1, MAP2, and TBR1). The sAD cortical neurons exhibited elevated total intraneuronal Aβ, thus successfully recapitulating a fundamental phenotype associated with sAD. Notably, our study revealed no observable morphological differences between the cortical neurons derived from control and sAD iPSCs despite higher intraneuronal Aβ levels in the sAD neurons. This profile is characteristic of early-stage pre-symptomatic models of AD and has been described in other studies [34–35]. In addition, elevated intraneuronal Aβ levels have been reported to be one of the earliest pathological events of AD, preceding both Aβ plaque accumulation and Tau tangles deposition, as well as synaptic dysfunction, suggesting a link between intraneuronal Aβ levels and early AD pathology [36–38]. Although the pathogenic role of elevated intraneuronal Aβ in early AD has been reported, its effect on proteasome proteolytic activity in early-stage sAD in human neurons has not been investigated.

Given the critical role of the proteasome in the maintenance of homeostasis in long-lived post-mitotic cells such as neurons, proteasome proteolytic activity may be a key factor in the etiology and pathogenesis of neurodegenerative disorders including sAD [14–15]. Thus, we hypothesized that an increase in intraneuronal Aβ levels will lead to an elevated proteasome activity as a compensatory mechanism at the early stage of sAD. Our analysis of the composition of active proteasomes revealed the robust expression of active 20S, 26S, and 30S proteasomes in both control and sAD neurons. To our knowledge, this is the first report showing the substantial expression of these proteasome assemblies in human iPSC-derived cortical neurons. While the composition of active 26S and 30S proteasomes revealed no significant differences between the control and sAD neurons, the composition of active free 20S proteasome showed a remarkable increase in the sAD cortical neurons. The 26S and 30S proteasome assemblies possess the 19RP on one or both ends respectively and are specialized for the targeted degradation of polyubiquitinated substrates in an ATP- and ubiquitin-dependent manner. However, the free 20S proteasome has no 19RP and is specialized for the degradation of substrates in an ATP- and ubiquitin-independent manner [11]. The differential expression of higher amount of free active 20S proteasome may suggest that the predominant proteasomal degradation in the early-stage AD cortical neurons does not depend on ATP, polyubiquitination, and binding to the 19RP. Compared to the 26S/30S proteasomes, the 20S proteasome is more stable and resistant to damage and has been found to exhibit high proteolytic activity during conditions of proteotoxic stress [39]. Because this response is conserved from yeast to humans [40], coupled with the fact that Aβ is a substrate of the human 20S proteasome [41], we conclude that the increased amount of active free 20S proteasome may be a compensatory response to the elevated intraneuronal Aβ in the sAD cortical neurons used as a human early-stage cellular model in this study. This conclusion is further supported by the recent finding of Liu et al. who reported increases in proteasome-related proteins in the cerebrospinal fluid of AD patients up to 20 years before the onset of clinical symptoms [42].

In the sAD iPSC-derived cortical neurons, the 20S proteasome exhibited increased proteolytic activity across all 3 proteolytic subunits when assessed using a single activity-based probe Me4BodipyFL-Ahx3Leu3VS that measures the 3 subunit activities simultaneously. The β5, β2, and β1 catalytic subunits (CT-L, T-L, and C-L activities respectively) are responsible for the total proteolytic activity of the 20S proteasome. The CT-L activity cleaves after amino acids with large or hydrophobic side chains, the T-L activity cleaves after basic residues, and the C-L activity is a post-glutamyl activity that recognizes acidic amino acid residues [43]. The β5-associated CT-L activity is deemed to be the initial and rate-limiting step in the proteolytic activity of the 20S proteasome. During this initial step, protein substrates are cleaved into large peptide fragments. In the following steps, the T-L and the C-L activities further break down the fragments generated in the initial step [44]. Beyond activity measurement, the protein expression of the β5 subunit of the 20S proteasome, also known as PSMB5, revealed no significant difference between the control and sAD neurons. These results imply that the proteasome subunits activation observed in the sAD iPSC-derived cortical neurons may involve subunit conformational changes or post-translational modifications that enhance the β5-associated CT-L activity without affecting subunit stability or expression. This might possibly be the case for the protein expression of the β1 and β2 subunits as well, however, we could not confirm this due to lack of specific antibodies for these subunits. The results of the activity-based probe were orthogonally supported by an extended independent kinetic analysis of the β5, β2, and β1 activities using distinct probes Suc-LLVY-AMC (β5), Boc-LRR-AMC (β2), and Z-LLE-AMC (β1) specific for each subunit activity under saturating conditions. The slope analysis of the kinetic assay curve revealed the β1, β2, and β5 proteolytic activities to have increased by 100%, 144.83%, and 55.75% respectively in the sAD neurons compared to the controls. The AUC analysis of the β1, β2, and β5 progress curves all revealed significant increases of 68.86%, 61.01%, and 113.90% respectively in the sAD neurons. Complementary use of the slope and AUC in this study allowed for the robust assessment of both early and late-phase kinetic profiles of the proteasome. A saturating condition implies that if the concentration of the substrate is sufficiently high in comparison to the concentration of the enzyme, then the rate of reaction will be proportional to the concentration of the enzyme. Thus, the amount of product formed in a given period of time would be proportional to the amount of active enzyme present, given that all other factors remain constant [45–47]. The 2.5µg total protein and 12.5µM substrates used in the study enabled us to achieve a saturating condition for the kinetic assay. As a result, the progress curves resulting from our kinetic assays of the β5, β2, and β1 proteolytic activities revealed a linear relationship supporting the establishment of a saturating condition between the proteasome subunits and their individual substrates.

The slope was estimated from the initial portion of the progress curve and represents the instantaneous rate of the initial linear phase of the proteasome-substrate reactions.

Because the slope captures the initial rate of the proteasome-substrate reaction, it is ideal for detecting early-phase changes in activity. Using the slope as a measure of proteasome proteolytic activity has been reported in other studies [48–49]. However, the proteasome is a major cellular degradative machinery and is continuously involved in proteolytic activities required to maintain intracellular homeostasis. Therefore, an instantaneous metric alone such as the slope may not be sufficient to robustly assess proteasome proteolytic activity under prolonged conditions as found intracellularly. In enzyme kinetics, fast-acting competitive modulators often influence the initial rates of the progress curve dramatically and are thus measured in the slope. However, slow-acting noncompetitive modulators tend to exert more pronounced effects on cumulative activity, especially if they gradually influence the activity of an enzyme over time [50–53]. This provides a compelling case for the analysis of the full progress curve rather than just the initial rates. By implication, a slow endogenous modulation of the proteasome is likely to be more accurately recorded by the AUC. This is because the AUC integrates the overall kinetic trace of the progress curve across the assay time, irrespective of linearity, to sum both the initial and late-phase proteolytic events. In this study, the combination of slope and AUC metrics allowed us to expand the dynamic range of our kinetic analysis to reflect both fast and sustained proteasome proteolytic activities. Notably, the slope and AUC of the proteasome proteolytic activities in our study validate each other, thus eliminating false positives/negatives.

In conclusion, our findings demonstrate the robust expression of active 30S, 26S, and 20S proteasome assemblies in human iPSC-derived cortical neurons. Using these neurons as cellular models of early-stage AD, we revealed that the level of active 20S proteasome and its catalytic subunits were more prominently expressed in sAD human iPSCs-derived cortical neurons despite increased intraneuronal Aβ. Our data suggest that increased proteasome activity might be a compensatory mechanism at the early stage of AD. Our findings support a central role for the proteasome in sAD pathogenesis and demonstrate the utility of iPSC-derived cortical neurons for investigating early-stage molecular underpinnings of the disease.

## Acknowledgement

This project has received funding from the European Union’s Horizon Europe research and innovation programme under HORIZON-MSCA-2021-PF-01 grant agreement No. 101069028 (to A.C.A) and the Sigrid Jusélius Foundation grant number 003701165704 (to J.K.). The UCI-ADRC is funded by NIH/NIA Grant P30 AG066519.

## Declaration of Generative AI and AI-assisted technologies

AI was not used in the preparation of the manuscript.

## CRediT authorship contribution statement

**Aderemi Caleb Aladeokin:** Writing – original draft. Writing – review & editing. Conceptualization. Methodology. Investigation. Formal analysis. Visualization. Validation. Data curation. Funding acquisition. Project administration. **Michael Jeltsch:** Writing – review & editing. Methodology. Validation. **Hayk Davtyan:** Methodology, Resources, Funding acquisition. **Mathew Blurton-Jones:** Methodology, Resources, Funding acquisition. **Jari Koistinaho:** Writing – review & editing, Methodology, Validation, Resources, Funding acquisition, Supervision.

