## Supplementary for "Composition and activity of the proteasome in human iPSC-derived neuronal model of early-stage sporadic Alzheimer’s disease"

### Methods

#### Maintenance and culture of human induced pluripotent stem cells

The human iPSC lines were generated by the University of California Irvine Alzheimer's Disease Research Center (ADRC) Induced Pluripotent Stem Cell Core (U.S.A), using non-integrating Sendai virus (Cytotune). The iPSCs were confirmed to be sterile and karyotypically normal via G-banding. Pluripotency of all lines was confirmed via (http://pluritest.org) and further confirmed using the Human Pluripotent Stem Cell Functional Identification Kit (R&D Systems), per manufacturer's instructions. Subject fibroblasts were collected and reprogrammed under approved IRB and human Stem Cell Research Oversight (hSCRO) committee protocols. Informed consent was received from all participants. Human iPSC line MAD 8 was generated as previously described [1]. All iPSC lines were maintained under feeder-free conditions using E8 medium (ThermoFisher Scientific) on plates coated with Matrigel (LDEV-Free, growth factor reduced; Corning, NY, USA) in a humidified incubator at 37°C with 5% $CO_2$. Cultures were fed fresh media daily and passaged every 3-4 days in small colonies using 0.5 mM EDTA (ThermoFisher

Scientific) and replated with 10μM ROCK inhibitor Y-27632 (Selleckchem) for the first 24 hours to enhance cell survival.

**Table S1:** Human iPSC lines used in this study

| iPSC line | Sample source | Donor age | Sex | Syndrome | APOE |
|---|---|---|---|---|---|
| ADRC 53 | Fibroblast | 76 | Female | Normal | E3/E3 |
| ADRC 54 | Fibroblast | 86 | Female | Normal | E2/E3 |
| MAD 8 | Fibroblast | 64 | Male | Normal | E3/E3 |
| ADRC 33 | Fibroblast | 73 | Female | AD | E3/E4 |
| ADRC 56 | Fibroblast | 79 | Male | AD | E3/E4 |
| ADRC 67 | Fibroblast | 85 | Male | AD | E3/E3 |

**Proteasome activity-based profiling with $Me_4$BodipyFL-$Ahx_3Leu_3$VS**

The activity-based profiling (ABP) of the 20S proteasome proteolytic subunits was performed using the fluorescent activity-based probe $Me_4$BodipyFL-$Ahx_3Leu_3$VS (UbiQ Bio). The probe allows a direct readout of proteasome activity at the individual subunit level by visualization of catalytically active 20S proteasome subunits via binding of its vinyl sulfone (VS) war head to the N-terminal catalytic threonine residue [2]. This cell-permeable probe specifically labels catalytically active proteasome β-subunits and allows for the detection of proteasome proteolytic activity in complex biological samples. Human iPSC-derived neurons were lysed in native lysis buffer (50 mM Tris-HCl pH 7.5, 150 mM NaCl, 5 mM $MgCl_2$, 0.5% NP-40, 1 mM ATP, 1 mM DTT) supplemented with protease and phosphatase inhibitors. Total protein concentration was determined using the BCA method (ThermoFisher). Equal amounts of protein (15μg) were incubated with 0.5μM of $Me_4$BodipyFL-$Ahx_3Leu_3$VS for 1 hour at 37°C. The incubation was stopped by adding 20% SDS to a final concentration of 1% SDS. Protein extracts were separated by SDS-PAGE using 4-20% Bis-tris gradient gel (GenScript) under reducing conditions. Fluorescently labeled proteasome subunits were visualized directly in the gel using G-Box (Syngene). Densitometric analysis was performed to quantify the relative abundance of active proteasome subunits as a percentage of the control by setting the control value to 100%.

**In-solution proteasome activity kinetic assays**

Briefly, cell lysates prepared in activity buffer (25 mM Tris, 20 mM KCl, 1 mM $MgCl_2$, 1 mM ATP, 0.5 mM DTT, and 0.05 mg/ml BSA) were normalized for total protein concentration and

2.5 μg of total protein were loaded into a black-walled 96-well plate with clear bottom. The reaction was started by adding a final substrate saturating concentration of 12.5 μM for Suc-LLVY-AMC (chymotrypsin-like activity, β5 subunit), or Boc-LRR-AMC (trypsin-like activity, β2 subunit), or Z-LLE-AMC (caspase-like activity, β1 subunit) in activity buffer, at the zero-time point. All samples were assayed with and without proteasome inhibitor epoxomicin or MG-132 to determine background AMC cleavage by other proteases in the samples. Plates were incubated in the dark for 10 minutes before measurement. Proteasome activity was determined for a single sample by subtracting the relative fluorescence of epoxomicin or MG-132 treated samples from that of the same sample not treated with inhibitor. Fluorescence accumulation upon proteasomal degradation of the fluorogenic substrates (360 nm excitation, 460 nm emission) was measured at 37°C on a Varioskan microplate fluorometer (ThermoFisher) every 1 min for 140 min. The kinetic approach continuously recorded the β5, β2, and β1 proteolytic activities over an extended period to produce real-time data, enhanced precision, and increased sensitivity for the identification of subtle activities that may be undetected in end-point single-timepoint activity assays. The slope of the linear portion of the progress curve and the area under the curve (AUC) were utilized as assay metrics. The combined use of the slope and AUC as assay metrics provided greater mechanistic insights into the individual proteolytic activities of the β1, β2, and β5 subunits and thus offers significant advantages over using either metric alone [3]. The slope, which captures how quickly an enzyme converts a substrate to product under optimum condition, was used to estimate proteasome efficiency per unit time [4]. The area under the curve (AUC), computed using the trapezoidal rule, which integrates the entire reaction over time, was used as a measure of the total proteasome activity across the progress curve [5-6].

**Results**

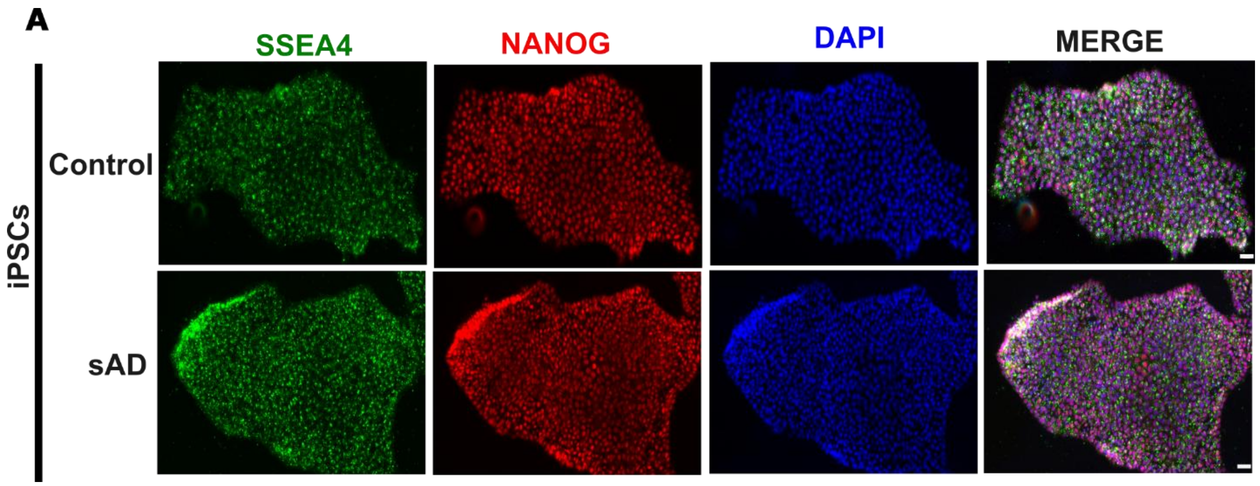


**Figure S1A:** Immunostaining of the pluripotency markers SSEA4 (green) and NANOG (red) in control and sAD iPSCs. DAPI was used to stain the nuclei. Scale bar = 20μm.

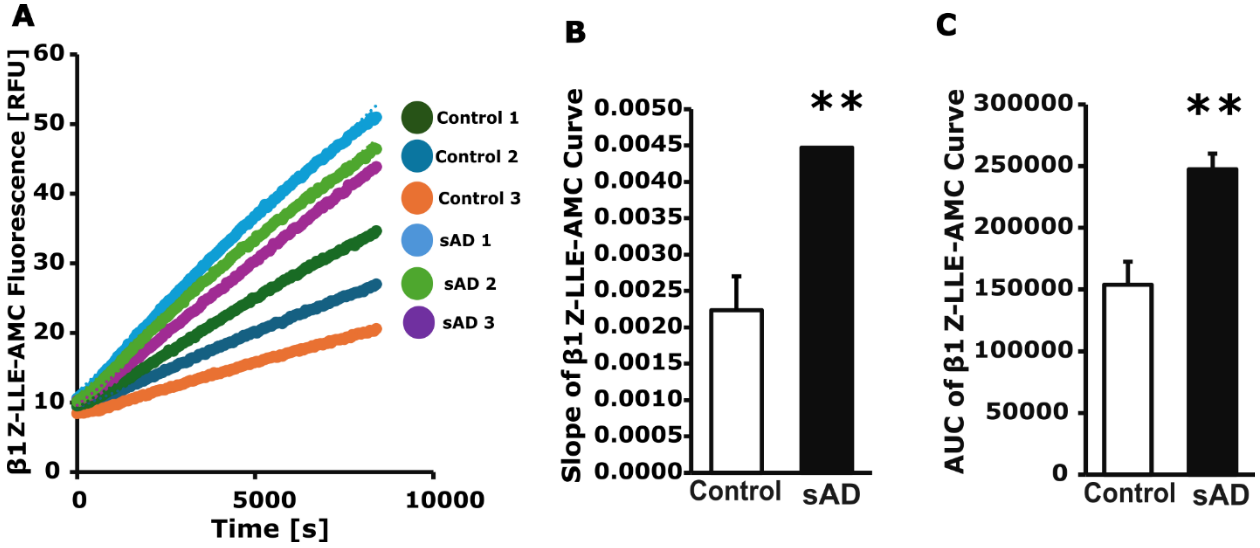

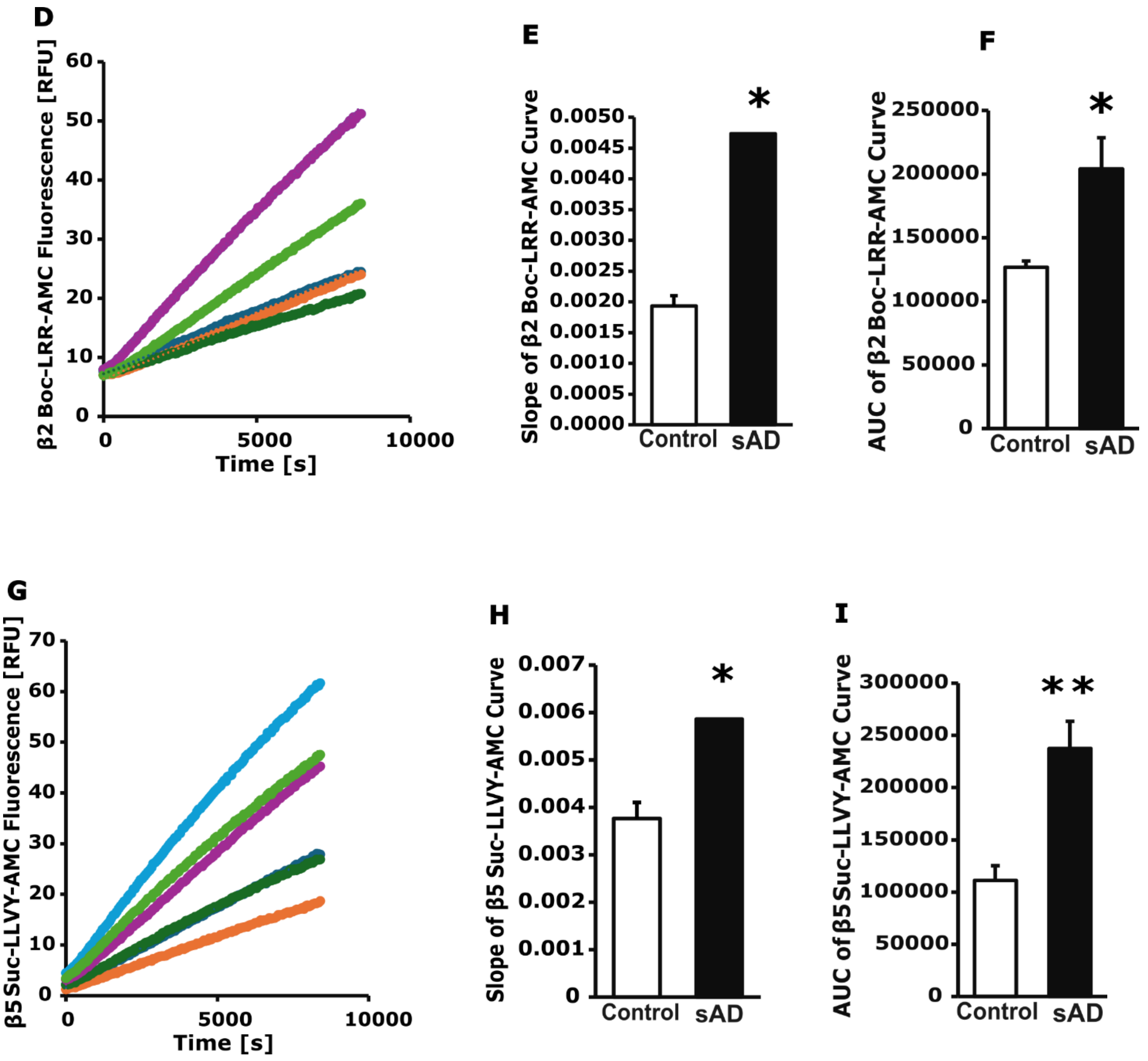


**Figure S6: Kinetic analysis of the proteolytic subunits β1, β2, and β5 of the 20S proteasome in control and sAD cortical neurons.** (A) Progress curve of β1 subunit activity showing the C-L activity measured using cleavage of Z-LLE-AMC peptide. (B-C) Slope ($p = 0.006$) and AUC ($p = 0.007$) of β1 subunit activity. (D) Progress curve of β2 subunit activity showing the T-L activity measured using cleavage of Boc-LRR-AMC peptide. (E-F) Slope ($p = 0.021$) and AUC ($p = 0.018$) of β2 subunit activity. (G) Progress curve of β5 subunit activity showing the CT-L activity measured using cleavage of Suc-LLVY-AMC peptide. (H-I) Slope ($p = 0.037$) and AUC ($p = 0.007$) of β5 subunit activity. Data represents mean ± SEM ($n = 3$).

Full gel of PSMB5

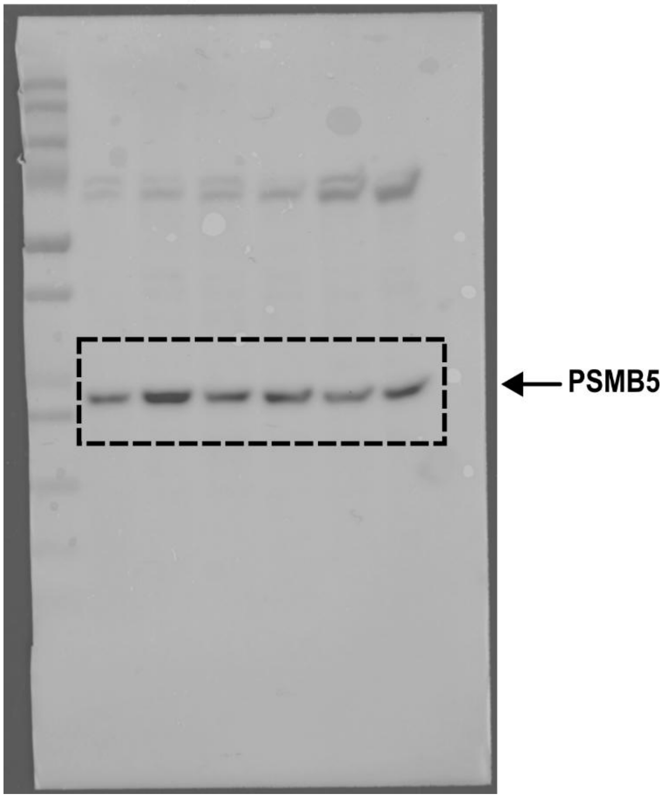


Full gel of GAPDH

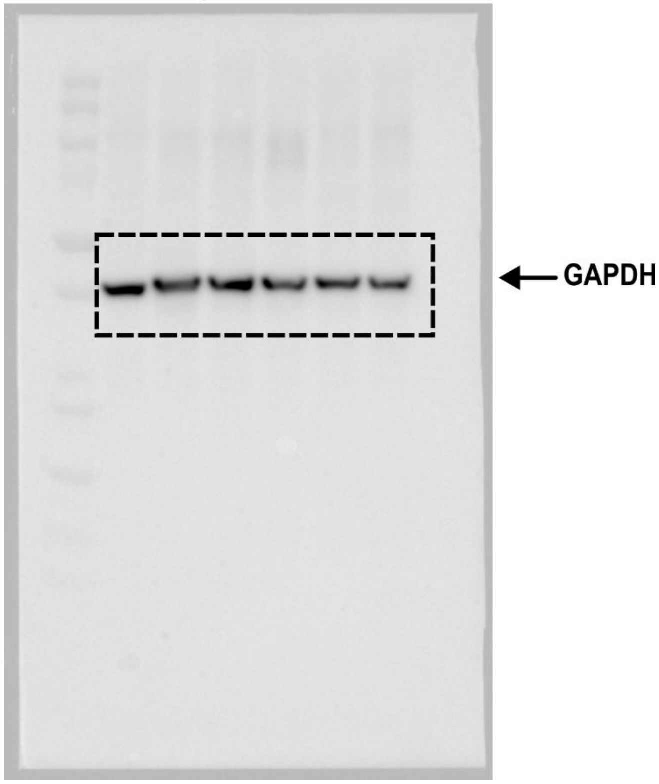
